# Gene-by-environmental modulation of longevity and weight gain in the murine BXD family

**DOI:** 10.1101/776559

**Authors:** Suheeta Roy, Maroun Bou Sleiman, Pooja Jha, Evan G. Williams, Jesse F. Ingels, Casey J. Chapman, Melinda S. McCarty, Michael Hook, Anna Sun, Wenyuan Zhao, Jinsong Huang, Sarah M. Neuner, Lynda A. Wilmott, Thomas M. Shapaker, Arthur G. Centeno, Khyobeni Mozhui, Megan K. Mulligan, Catherine C. Kaczorowski, Liza Makowski, Lu Lu, Robert W. Read, Saunak Sen, Richard A. Miller, Johan Auwerx, Robert W. Williams

**Affiliations:** Department of Genetics, Genomics and Informatics, University of Tennessee Health Science Center, Memphis, Tennessee 38163, United States; Department of Preventive Medicine, University of Tennessee Health Science Center, Memphis, Tennessee 38163, United States; Department of Medicine, College of Medicine University of Tennessee Health Science Center, Memphis, Tennessee 38163, United States; Laboratory of Integrative Systems Physiology, Interfaculty Institute of Bioengineering, École Polytechnique Fédérale de Lausanne (EPFL), Lausanne CH-1015, Switzerland; Department of Biology, Institute of Molecular Systems Biology, ETH Zurich, Zurich CH-8093, Switzerland; Department of Pathology, University of Michigan Geriatrics Center, Ann Arbor, Michigan 48109, United States

**Keywords:** Longevity, lifespan, high fat diet, body weight, genetic reference population, gene by environment (GXE)

## Abstract

Diet and environment profoundly modulate lifespan. We measured longevity as a function of diet and weight gain across a genetically diverse family of mice. We followed 1348 females from two parental strains—C57BL/6J and DBA/2J—and 146 cohorts of BXD isogenic progeny strains (*n* = 73) across their lifespan on a low fat chow diet (CD, 18% calories from fat) and on a high fat diet (HFD, 60% calories from fat). On average, HFD shortens lifespan by 85 days or 12%, roughly equivalent to an 8–10 year decrease in humans. However, strain variation in the response of diet on longevity is remarkably high, ranging from a longevity loss of 54% in BXD65 to a gain of 37% in BXD8. Baseline weights and early weight gain are both associated with a mean decrease in longevity of ∼4 days/g. By 500 days-of-age, cases fed HFD gained four times as much weight as control on average. However, strain-specific variation was substantial, thus weight gain did not correlate well with lifespan. In summary, high fat had a strong negative effect on longevity, but genetic interactions effects were even stronger. This highlights the unequivocal importance of genetic differences in making dietary recommendations.

## Introduction

Longevity is among the most heterogeneous and complex of traits. Differences in lifespan are dependent on innumerable gene-by-environmental (GXE) interactions (de Magalhães et al., 2012, Kuningas et al., 2008, McDaid et al., 2017, Hook et al., 2018). Nutrition, in particular, has a profound influence on health and lifespan (reviewed by Fontana and Partridge, 2015). Relative to an ad libitum diet, caloric restriction and intermittent fasting improve lifespan and health (reviewed by Heilbronn and Ravussin, 2003; Liang et al., 2018; Speakman et al., 2016). Effects of restricted diets and intermittent fasting are not entirely dependent on patterns of caloric intake, but depend on dietary macro- and micronutrient composition, the amount of time spent in different metabolic states, age of onset, periodicity of access to food, sex, and perhaps of most importance to us, differences in genotype (Vaughan et al., 2018) and to gene-by-diet interactions (Barrington et al., 2018).

The mouse is an excellent model for research at the interface of metabolism and aging, sharing most protein-coding genes with humans (Pennacchio and Rubin, 2003), but with a much shorter life span that enables longevity studies in controlled environments and under multiple experimental and dietary conditions (Miller et al., 2007, Yuan et al., 2011, Strong et al., 2013). However, most studies using animal models unfortunately do not incorporate a level of genetic complexity that matches typical human populations (Saul et al., 2019; Williams, 2006; Williams and Williams, 2017). Effects of DNA variants and dietary, drug, or environmental perturbations are often studied on a single genome—often that of the inbred C57BL/6 strain. This intense single-genome focus compromises translational utility of discoveries.

To address this problem, we rely on a very large family of BXD strains of mice (*n* = 150) that segregate for over 6 million variants (Peirce et al., 2004, Wang et al., 2016, Ashbrook et al., 2019). Collectively, the family also incorporates an impressive level of variation in phenotypes related to aging, metabolism, and mRNA and protein expression in liver, muscle, brain, and many other tissues and cell types (Williams et al., 2014, Houtkooper et al., 2013, Andreux et al., 2012, Houtkooper et al., 2011, Gelman et al., 1988, De Haan and Van Zant, 1999).

Previous studies of about 30 strains of the BXD family (Gelman et al., 1988; Lang et al., 2010) demonstrate at least two-fold variation in lifespan on a standard diet—from approximately 12–15 months for the shortest-lived strains to 30 months for the longest-lived. In these studies, conventional heritabilities of lifespan are as high as 25–45%, but the effective heritabilities (*h*^*2*^_*RI*_) which properly account for the depth of resampling (*n* = 8 to 12 replicates/genome) are as high as 80% (Belknap, 1998; Hook et al., 2018). The BXD family is particularly well suited to study GXE interactions because diverse but perfectly matched cohorts can be treated in parallel using different diets (Rikke et al., 2010, Hall et al., 2014, Andreux et al., 2012, Williams et al., 2016; Wu et al., 2014). The effect of genetic variation has been well studied in the context of dietary composition and caloric restriction on lifespan (Finkel, 2015, Keipert et al., 2011; Skorupa et al., 2008). However, key results remain controversial. While caloric restriction is undoubtedly advantageous in boosting longevity *on average*, there is good evidence that effects are not universal, and that certain individuals and genometypes do not benefit in all environments (Barrington et al., 2018; Liao et al., 2010; Mitchell et al., 2016; Rikke et al., 2010).

In this study, we have measured longevity, body weight, and weight gain across two cohorts of fully sequenced, and highly diverse strains (Pierce et al., 2004; Ashbrook et al., 2019). We studied females in a tightly controlled environment on diets that differed greatly in fat content—those on a standard low-fat chow diet (CD, 18% of calories from fat) and those on a very high-fat diet (HFD, 60% cal from fat).

To the best of our knowledge this is the largest GXE experiment on the effects of a high fat diet on longevity and weight change, and includes matched data for 1348 cases, two cohorts, and 73 BXD genometypes.

We address the following questions:

1. What is the average impact of a high fat diet that is otherwise matched for protein content, on longevity across members of the BXD family? Does the HFD influence the level of variability in longevity?
2. To what extent do genetic differences among strains modulate effects of HFD relative to CD? Put another way: What is the strength of evidence in favor of GXE effects on longevity?
3. Does baseline body weight at young adulthood (∼120 days age) predict longevity, and is the change in body weight in response to HFD a strong predictor of longevity?
4. To what extent is weight gain *per se* linked to a reduction in longevity, and how does weight gain vary among strains on high and low-fat diets?

## Methods

### Animals and Diets

Animals were raised and housed in a specific pathogen-free (SPF) facility at UTHSC (Memphis, TN), at 20–24 °C in temperature on a 12-hour light cycle. During the course of our study, serum samples from sentinel mice were tested quarterly for the following pathogens—ectromelia virus, epizootic diarrhea of infant mice (EDIM), lymphocytic choriomeningitis (LCM), *Mycoplasma pulmonis*, mouse hepatitis virus (MHV), murine norovirus (MNV), mouse parvovirus (MPV), minute virus of mice (MVM), pneumonia virus of Mice (PVM), respiratory enteric virus III (REO3), Sendai, and Theiler’s murine encephalomyelitis (TMEV GDVII). Semi-annual necropsies are performed to test for endoparasites by microscopic examination of intestinal contents and anal tape preparations and ectoparasites by direct pelt microscopic examination. All such tests were negative throughout the study.

From October 2011 through to December 2018, animals from both parental strains, C57BL/6J and DBA/2J, and 73 BXD strains were followed from their entry into the aging colony from a large breeding colony — typically around 120 ± 66 days of age but with a wide range, from 26 days to 358 days—until their death. Animals were inspected daily, and deaths were recorded for each animal with a precision of one day. Moribund animals (∼10%) were euthanized, and those above the age of 200 days were included in longevity calculations. Criteria for euthanasia were based on an assessment by our veterinary staff following AAALAC guidelines. All animals were initially raised by dams on the standard chow diet in the breeding colony. Upon entry into the aging colony, females were aged in groups of up to 10 in polypropylene cages (935 cm^2^) provisioned with Envigo Teklad 7087 soft cob bedding. Animals were provided either the standard low fat chow diet (CD, Envigo Teklad Global 2018, 18.6% protein, 18% calories from fat (fatty acids comprise of 0.9% saturated, 1.3% monounsaturated, and 3.4% polyunsaturated fats), caloric content of 3.1 kcal/g), or a high fat diet (HFD, Envigo Teklad TD06414, 23.5% protein, 60.3% calories from fat (fatty acid profile comprises of 37% saturated, 47% monounsaturated, 16% polyunsaturated fats), caloric content of 5.1 kcal/g). For detailed diet data sheets see Supplemental Information. Animals had *ad libitum* access to food and aquifer-sourced autoclaved municipal tap water.

We studied a total of 1348 individuals (*n* = 663 on CD, *n* = 685 on HFD). Animals were labeled using ear tags, and were randomly assigned to a diet. Baseline weight was measured at entry into the study. Seventy-seven percent (*n* = 527) of animals started on HFD at ages between 50–185 days, but some started on the diet at ages as early as 26 days or as late as 358 days. Fewer than 2% of animals (*n* = 12) were placed on HFD at an age greater than 365 days, and these were not included in the analysis. Less than 20% of animals were retired breeders that entered the study at 180+ days of age. Each animal was weighed to the nearest 0.1 gram every other month from start of diet until death (Table S2). A separate subpopulation of 662 animals (*n* = 333 on CD, *n* = 329 on HFD) from matching BXD strains were sacrificed at specific time-points (approximately 6, 12, 18 and 24 months) for tissue collection across both cohorts (Table S3, Williams EG et al., manuscript in preparation). Organ weight data at 18 months-of-age from these animals were included in the analysis for this study. The aging colony at UTHSC is still in operation, but for this analysis we only consider animals with deaths between April 2012 and November 2018.

The colony was moved to a new vivarium in the Translational Science Research Building in April 2016 from the Nash Annex Building, both at UTHSC. Approximately 60% of the animals lived and died in the original Nash Annex vivarium, ∼35% were born in the Nash Annex but lived in both vivaria, and ∼5% were born and spent their entire lives in the new facility. We carefully evaluated birth and death data over all seasons from both vivaria to successfully rule out any site-specific or seasonal effect on longevity.

Most animals were fixed by immersion in 10% neutral buffered formalin within 24 hours of death. The body cavity was opened prior to immersion to improve preservation. Evenly balanced cohorts on the diets were selected based on fixation quality for necropsy with histopathology of tissues. A board-certified veterinary pathologist (RWR) performed necropsies and judged probable cause of death and other morbidities.

All experimental procedures were in accordance with the Guidelines for the Care and Use of Laboratory Animals published by the National Institutes of Health and were approved by the UTHSC institutional Animal Care and Use Committee.

### GeneNetwork

Mean, median, and 75% quantile longevity data from both cohorts are available in GeneNetwork2. https://www.genenetwork.org (GN2) under the headings **Species:** *Mouse (mm10)*; **Group:** *BXD Family*; and **Type**: *Phenotypes* (e.g., GN2 traits BXD_18435, 18441, 19451, 19452, 21302, 21450). Body weight data at 6, 12, 18 and 24 months is also available in GN2 (e.g., traits BXD_19126, 19130, 19131, 19167, 19168, 19169, 19170, and 19171). Organ weight data on both diets, including liver, heart, kidneys and brain, at 18 months can be retrieved, but is only covered briefly in this report (traits BXD_20156, 20157, 20158, 20159, 20353, 20354, 20148, 20149, 20150, 20151, 20146, 20147). Finally, individual data are also available for all cases, both as Supplementary tables (Table S1, S3) and in GN2 under the headings **Species:** *Mouse (mm10)*; **Group:** *BXD NIA Longevity Study*; and **Type**: *BXD-NIA-Longevity Phenotypes*.

### Statistics

In some respects, some aspects of our design are similar to an observational prospective study, rather than a rigidly controlled experiment—the main factor being that cases were place on the HFD over a range of ages (see Results for details). We used statistical methods that are common to observational data analysis to adjust for variability in age of entry.

Longevity and body weight data were stratified by diet and by strain. Effects of diet, strain, and body weight on longevity were analyzed using a random-effects model in R using the *metafor* package (Viechtbauer, 2010) and a mixed-effects Cox proportional hazard model using Therneau’s *coxme* R package 2.2.-10 (*cran.r-project.org/web/packages/coxme/index.html*) (Therneau and Grambsch, 2000). Survival analyses were performed using the survival package for R and the data were right-censored (e.g. Fig. 2A-C, censored cases CD, *n* = 32; HFD, *n* = 80). Survival curves were computed by ANOVA and regression analyses were performed using R. Results were also tested using Wald test, likelihood ratio test, and Wilcoxon test. All graphs were made in R, and final figures were all prepared with Adobe Illustrator.

## Results

### High fat diet shortens lifespan but with considerable variation among strains

Balanced sets of 1348 females from 73 strains were assigned to either CD (*n* = 663) or HFD (*n* = 685) at an average of 120 days of age, weighed every other month and followed until natural death (Figure 1A). The HFD has a strong average effect on lifespan across the family. Mean lifespan decreased significantly (*p* <2.2E-16, *r* = 0.2) from 690 ± 8 SE (±199 SD) days on the control diet to 605 ± 6 SE (±169 SD) days on the HFD. Median longevity decreased 77 days—from 703 to 626 (Figures 1B, box plot inset). The 75% quantile change in longevity decreased from 796 days to 699 days. Assuming linear scaling and that a dietary change was imposed in humans at about 20 years-of-age, this 77-to 97-day difference would compare roughly to a 8-10 year loss of longevity in humans (Flurkey K, Currer JM, Harrison DE. 2007).

**Figure 1.**
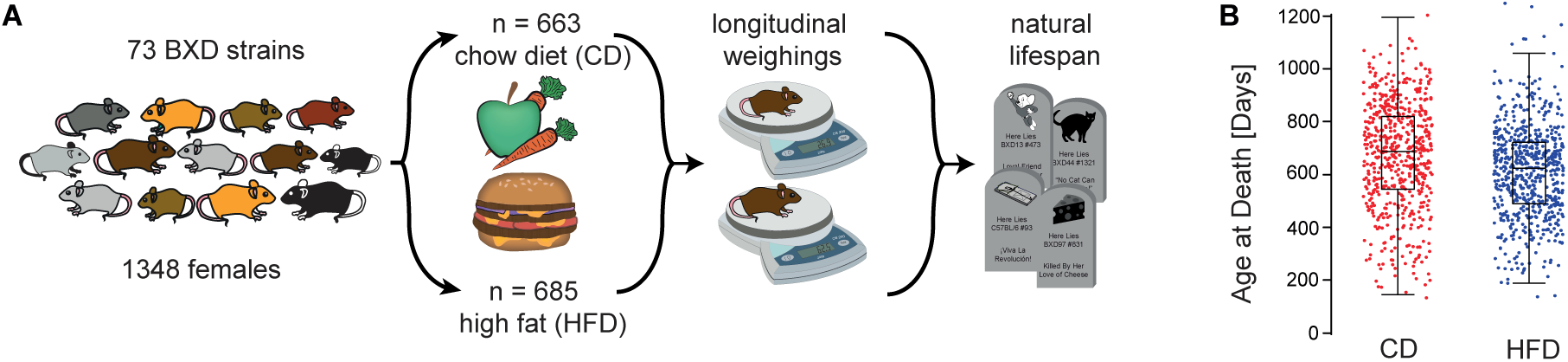
Study Overview (**A**) Balanced sets of females from 73 BXD strains were assigned to the low fat chow diet (CD) or the high fat (HFD) diet and weighed every other month. (**B**) High fat diet modulates longevity. When grouped by diet—irrespective of genetic background—median lifespan of CD cohort exceeded by 77 days that of the HFD cohort (see *box plot* inset). **Red** and **blue** dots represent individual cases on CD and HFD. For strain averages of longevity in the two cohorts see Supplementary figure **S1**, panels **A** and **B**.

**Figure 2.**
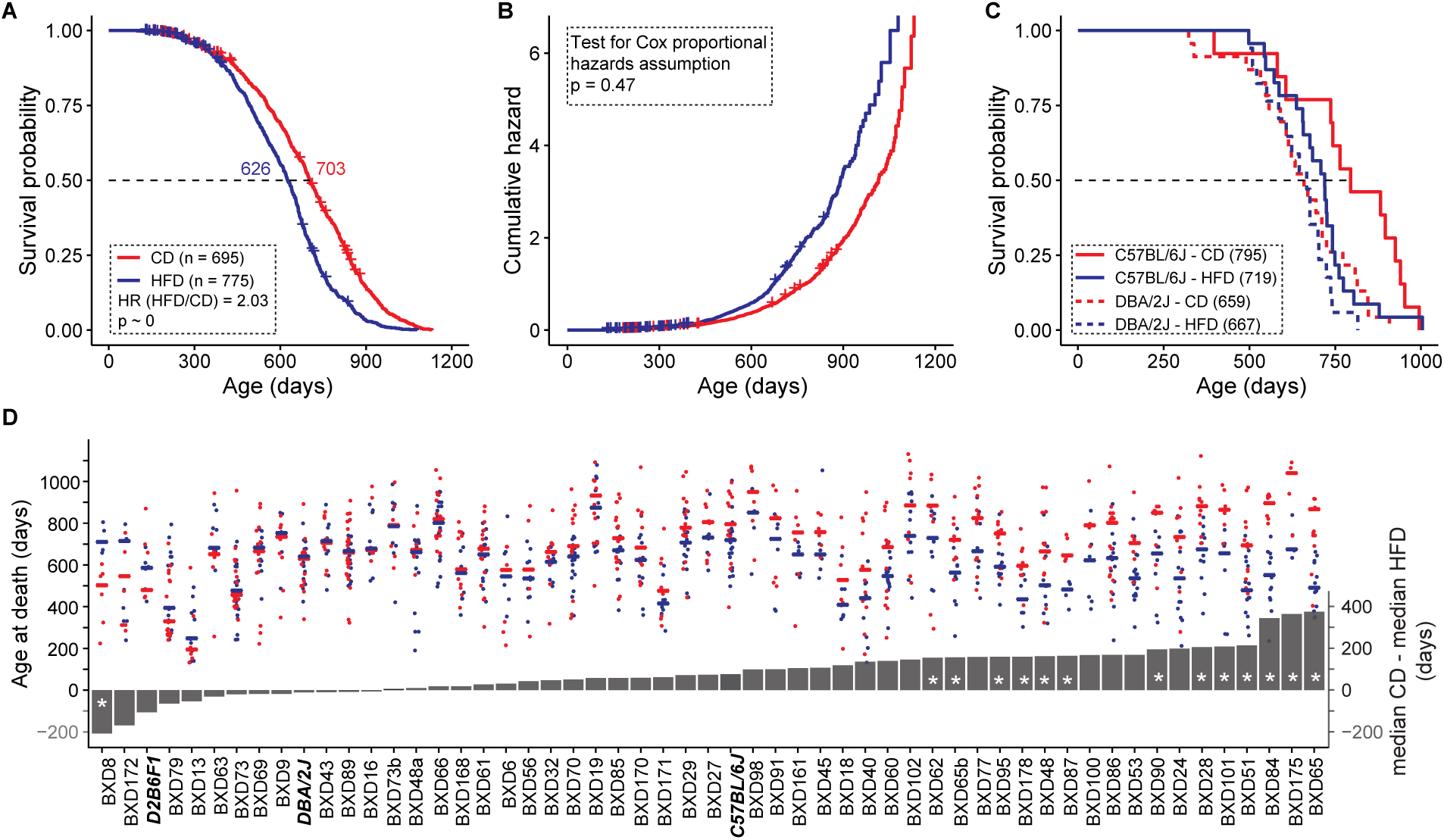
Diet influence on longevity (**A**) Median longevity decreased by 77 days on HFD. Cases fed HFD have a two-fold higher risk of death compared to those fed CD. (**B**) Cumulative hazard curves by diet do not cross and the hazards ratio of 2.0 is relatively constant throughout the study. (**C**) The lifespan of C57BL/6J, but not DBA/2J, is influenced by diet. **Numbers in parentheses** are median lifespans in days. (**D**) Longevity on a high fat diet depends strongly on strain. **Red** points represent longevities of cases on CD and **blue** points those on HFD. Lines represent median survival (**left y axis**). Grey bars represent the difference in median survival on the diets (**right y axis**). Negative values to the left indicate higher survival on HFD, positive values indicate higher survival on CD. Parental strains and F1 are denoted by **bold italics font. Asterisks** in bars denote significant FDR scores at a q value of 0.1. Censored cases in A–B are still alive and are marked by **+** signs.

Using a mixed-effects Cox model with diet as a fixed effect and strain as random effect, we estimated a hazard ratio of 2.0, indicating that animals on HFD have two-fold higher age-adjusted risk of death compared to their perfectly genotype matched CD-fed controls (Figure 2A). The hazard ratio is relatively constant throughout the study and there is no crossing of the cumulative hazard curves (Figure 2B, *p* = 0.47).

While HFD decreased lifespan at the family level, individual strains differ greatly in how they react to diet (Supplemental Figure S1 A-B). The parents exemplify the difference—the longevity of the DBA/2J paternal strain in unaffected by diet, whereas the C57BL/6J maternal strain loses 76 days on the HFD (Figure 2C). In fact, some progeny even lived longer on the HFD, demonstrating that the longevity-diet relation is modulated by a forceful GXE component (Figure 2D). Overall, longevity in 21 strains out of 67 (strains with ≥ 4 natural deaths in both cohorts) was significantly affected by HFD in either direction at a nominal *p* threshold of 0.05. To correct for multiple testing (*n* = 67) we computed *q* values (see Figure 2D, asterisk**\*** symbol for *q* value < 0.1) and with this correction, only 15 strains have significantly different longevity. Interestingly, BXD8 has significantly improved longevity on a high fat diet, with a median increase in longevity of 208 days (*t* = 4.0, *p* = 0.0052, *q* <0.05, two-tailed), with one other strain trending in the same direction—BXD172 (146 day increase on HFD, *p* = 0.073), but with a high *q* value of 0.85. As expected, high fat reduced longevity for most strains (Fig 2D, right side)—most prominently for BXD65, with a decrease in median longevity of nearly a year (345 days, *t* = 9.3 *p* = 9.0E-7, *q* = 6.0E-7). Genetic variance explains 30% of the total variance in lifespan while diet accounts for 5%, and gene-diet interactions account for 7%.

### Does a high fat diet modulate variability in longevity?

We computed coefficients of variation (CV, SD/mean) for all strains in the two cohorts with sample sizes of greater than 6 cases (GN2 traits BXD_21533 and BXD_21534, Supplementary Figure S2). The mean CV on the control diet is 0.27 ± 0.015 SE (*n* = 52) compared to 0.25 ± 0.013 SE on HFD (*n* = 50)—a small drop in CV on the HFD, but not significant. Ranges of CV among strains are high—from a low of about 0.1 to a high of 0.5. But only 10% of strains have CV above 0.4, while 10% have CVs below 0.15. Much of this range is presumably sampling error. The CVs of the inbred strains are slightly higher on average that those of the outbred female mice of about 0.21 (Miller et al., 2007, see GN2 Trait ITP_10001).

### Body weight at young adulthood is a strong predictor of longevity

Cases were weighed regularly throughout their lifespan. As expected, those on HFD gained more weight on average than those on CD (Figure 3A). Initial weights at entry into the aging colony at about 120 days of age had a significant influence on lifespan after adjusting for differences in baseline age (*p* <0.001, *r* = 0.1) (Figure 3B). A one-gram increase in weight at this stage is associated with a 5-day loss of longevity. By design there is no appreciable difference between cases assigned to the HFD (23.11 ± 0.22 SE g, *n* = 685) versus those continuing on the CD for the remainder of their lives (23.26 ± 0.22 g, *n* = 659). Of interest, the slope of –5 days/g at ∼120 days is not affected by the subsequent diet. This is highlighted by the parallel red and blue lines in Figure 3B, and indicates that the young adult body size-longevity relation is relatively insensitive to GXE.

**Figure 3.**
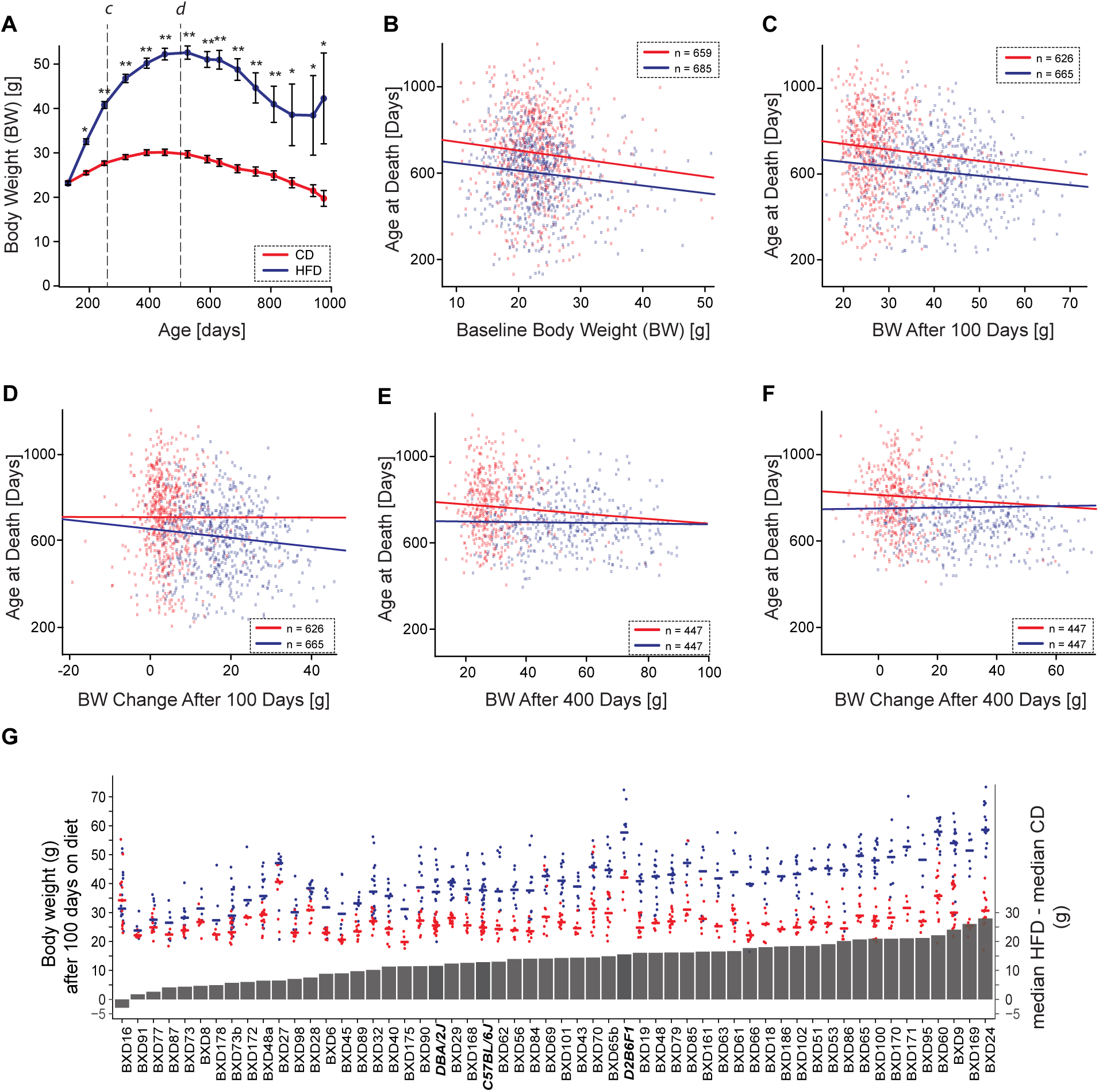
Effect of body weight on longevity (**A**) Body weight by diets and age (CD, **red** line and HFD, **blue** line). Single and double **asterisks** denote significance at *p* <0.05 and <0.001. Body weight declines on both diets after about 500 days of age. (**B**) Initial body weight—the weight at entry into colony prior to the point of being of any animals being shifted to the HFD—has a modest but consistent negative slope with longevity (–5 days/g, *p*<0.001, *r* = 0.1) that is not exacerbated by the HFD. (**C**) Body weight after 100 days on the two diets (∼260 days age) correlates negatively with longevity (–4 days/g, *p* <0.001, *r* = 0.3, see line labeled **c** in Panel **A**). (**D**) Early weight change in response to HFD (**blue** line)—the difference from baseline after 100 days on diet—was negatively related with longevity (–4 days/g, *p* = 0.004, *r* = 0.1), but this is not true of cases remaining on CD. (**E**) After 400 days on diet (∼500 days age), body weight does not predict variance in longevity (see line labeled **d** in Panel **A**) (*p* = 0.87, *r* = 0.01). (**F**) Substantial weight change after prolonged HFD feeding— difference from baseline to 400 days on diet (blue line)—does not predict of longevity (*p* = 0.65, *r* = 0.02). (**G**) Strain-wise changes in median weight after 100 days on diets. **Red** points represent longevity of cases on CD and **blue** points those on HFD. **Lines** represent median body weight (**left y axi**s). **Grey** bars represent differences in median body weight on the diets (**right y axis**). Parental strains and F1 hybrids are denoted by **bold italics font**. Strain averages of body weight at 500 days of age in the two cohorts are shown in see Supplementary figure **S1**, panels **C** and **D**.

### Early body weight gain is associated with a reduction in longevity

Body weight measured after 100 days on both diets also correlates negatively with longevity (Figure 3C), a one gram increase now corresponding to a decrease of 4 days (*r* = 0.3). This effect is observed even after adjusting for strain differences. Looking at change in body weight after 100 days on diet, early body weight gain in response to HFD, but not CD, is negatively correlated with longevity (*p* = 0.004, *r* = 0.1) (Figure 3D).

### Diet significantly alters longevity after adjusting for weight gain

We chose to focus on two time points for body weight analyses—100 days on diet as a point to evaluate initial weight gain on HFD, and 400 days on diet, a stage that is close to the maximal weight on both diets. The mean weight of the population plateaus around 500 days of age and declines thereafter on both diets. By 500 days of age, cases had been on HFD for 400 ± 44 days and gained an average of 29.5 g. Those on the CD gained only 6.2 g (mean weight on CD = 29.7 ± 0.35 SE g, *n* = 447; mean weight on HFD = 52.6 ± 0.63 SE g, *n* = 447).

Surprisingly, the substantial increase in body weight on the HFD does not significantly correlate with lifespan (Figure 3E). Only10% of the effect of diet on longevity is mediated through body weight gain after adjusting for strain-specific differences in longevity. Mirroring this interesting observation, sustained weight gain after prolonged feeding of HFD (Figure 3F) has no predictive value, emphasizing that the diet itself, rather than weight gain per se, modulates longevity. Overall, 53 out of 67 strains have significant weight gain on the HFD after 100 days (*p* <0.05, *q* <0.1, *t* values ranging from –3.03 to – 13.43) (Figure 3G). At 500 days of age, 45 out of 57 strains have gained significant weight on the HFD over 400 days (*p* <0.05, *q* <0.1, *t* values ranging from –3.03 to –13.14) (Supplemental Figure S1 C-D). Diet accounts for 45% of variance in body weight gain versus 22% explained by genotype, 11% by gene-diet interaction, and 22% as unexplained residual.

### Insufficient evidence of association between longevity and major metabolic organ weight

We compared organ and tissue weights of a separate subsample of animals on the diets at ∼500 days of age. HFD mice (*n* = 63) had 84% greater fat mass, 25% greater heart mass, 19% greater liver mass, and 18% greater kidney mass at ∼500 days compared to controls (*n* = 71). However, HFD did not influence brain mass. The correlations between longevity and organ weights at 18 months is not significant (Supplemental Figure S3 A-F).

### Major morbidities contributing to death in the aging colony

We carried out gross necropsy, with or without histopathology, for a total of 155 cases—76 animals (69 with a single cause of morbidity and 7 with multiple causes) from 45 strains on CD, and 79 animals (71 with a single cause of morbidity and 8 with multiple causes) from 43 strains on HFD. Likely cause of death was clear at necropsy or following histopathology in 87% of cases. Hematopoietic neoplasia—lymphomas and histiocytic sarcomas—was a leading cause of death, accounting for ∼35% of deaths in both cohorts. Miscellaneous non-neoplastic conditions causing significant morbidity or death were detected in 24% of CD and 32% of HFD-fed animals.

Preliminary evidence points towards the influence of diet on causes of morbidity and mortality (Table 1). In the HFD cohort there is an increased prevalence and severity of cardiovascular disease and lesions, including atrial thrombosis, cardiomyopathy and cardiac dilation sometimes with hepatic steatosis and centrilobular atrophy. Nineteen percent of these cases (15 out of 79) had heart pathologies rated moderate to severe, whereas only 1 of 76 CD cases had a heart lesion rated at least moderately severe (2-tail Fisher’s exact test *p* = 0.0003). Four HFD cases, but no CD cases, had systemic polyarteritis. Conversely, some pathologies are evidently more pervasive in the CD cohort. Thirty-six percent of CD mice displayed non-hematopoietic malignant neoplasia (27 of 76 CD–11 sarcomas, 15 carcinomas, and 1 teratoma) compared to 13% HFD mice with non-hemopoietic malignancy (10 of 79 HFD, 6 sarcomas and 4 carcinomas). While this preliminary finding is nominally significant (Fisher *p* = 0.0012), we have not corrected for multiple comparisons.

**Table 1.** Major morbidities contributing to death in BXD family.

| Morbidity Type | Diet | n with single morbidity | n with multiple morbidities | p-value (two-tailed Fisher's exact test) |
| --- | --- | --- | --- | --- |
| Unnatural causes | CD | 7 | 1 | 0.1257 |
|  | HFD | 3 | 0 |  |
| Non-neoplastic conditions | CD | 16 | 2 | 0.3715 |
|  | HFD | 22 | 2 |  |
| Heart/cardiovascular | CD | 0 | 1 | 0.0003* |
|  | HFD | 8 | 7 |  |
| Polyarteritis (autoimmune) | CD | 0 | 0 | 0.1204 |
|  | HFD | 3 | 1 |  |
| Lymphoma | CD | 25 | 3 | 0.7402 |
|  | HFD | 26 | 1 |  |
| Sarcoma | CD | 9 | 2 | 0.2042 |
|  | HFD | 4 | 2 |  |
| Carcinoma | CD | 12 | 3 | 0.0065* |
|  | HFD | 4 | 0 |  |
| Renal | CD | 0 | 1 | 0.3673 |
|  | HFD | 1 | 3 |  |
| Teratoma | CD | 0 | 1 | 0.4903 |
|  | HFD | 0 | 0 |  |

Unnatural causes = flooded cage, foot injury in wire lid, eye abrasion leading to euthanasia etc.
Miscellaneous non- neoplastic conditions = inflammation, infection, amyloidosis, ulcerative dermatitis
Heart = major contribution of cardiovascular failure other than polyarteritis
Polyarteritis = systemic multifocal arteritis suggestive of autoimmune disease
Lymphoma = hematopoietic neoplasia including lymphoma and histiocytic sarcoma
Sarcoma = nonhistiocytic sarcoma, spindle cell, hemangiosarcoma, leiomyosarcoma, sarcoma NOS
Carcinoma = carcinomas (hepatocellular, squamous cell)
Renal = renal failure due to severe nephropathy

## Discussion

### Synopsis

We have studied effects of a high fat diet on aging and lifespan in a large and genetically diverse family of mice. Our focus here is on the overall and genotype-specific effects of diet on longevity and weight gain, rather than on the genetic control of longevity or weight *per se*—work that will soon follow. In the introduction, we pose four sets of questions for which we now have good answers:

1. Yes, as expected a diet that is high in fat reduces longevity by an average of 12% in our specific cohorts, and as in humans, the risk of cardiovascular disease and cancers is elevated on the HFD.
2. However, this is not a universal response, and GXE effects are strong for both longevity and weight gain. Even after we correct for multiple comparisons, one strain lives significantly longer and another strain gains no weight on the HFD.
3. We confirm that lower weight at an early age is linked to longer longevity, and that this effect is also true on the HFD.
4. There is at best only a modest link between weight gain after maturity and longevity. Weight gain measured late in life—between ages of 12–18 months—accounts for minimal variation in longevity. And finally, diet modulates longevity and has a stronger effect on longevity than weight gain itself. By far the strongest effect on longevity and weight gain is the genometype of the animal.

In the interest of avoiding overly broad generalization from this study of GXE effects, we highlight two key design limits. First, we only studied females—the main reason being that unlike males, they can be housed without overt and detrimental aggression. Males generally do not live as long as females (Austad and Fischer, 2016), and in the large ITP study the overall difference is 10%—808 ± 2.88 days for males (*n* = 7147), 894 ± 2.43 days for females (*n* = 6139) (see GN2 Trait ITP_10001). Of even greater practical importance to us, variability of longevity is much higher in males than females—a coefficient of variation (CV) of 0.30 versus 0.21. As a result, studies of longevity in females are statistically twice as efficient as those using males (the ratio of the squares of CVs) (Cheng et al., 2019; Miller et al., 2007). We can only presume that the same GXE effects of diet on longevity apply to males (Andreux et al., 2012; Williams et al., 2016). Second, we have studied only one dietary contrast. Each diet has the potential to reveal novel GXE effects as a function of macro- and micro-nutrients and many other environmental factors (Barrington et al., 2018). One of the significant advantages of using cohorts of isogenic cases is that it become practical to build empirical maps of *reaction norms* for diet, health, and longevity, as done here for two diets (Griffiths et al., 2000). This highlights the suspect nature of *one-size-fits-all* dietary recommendations for highly diverse humans with equally diverse exposures and environments.

### GXE and longevity

A diet rich in saturated fats shortens lifespan by an average of almost three months in BXD females, roughly scaling to a decade decrease in humans. This diet is associated with an average two-fold higher age-adjusted risk of death compared to chow-fed controls. Longevity under the two diets correlates moderately (*r* = 0.60, GN traits BXD_18435 and 18441). However, strains display remarkably wide variation in responses to diet, and despite the strong effect, diet accounts for just 5% of the total variance in longevity. In comparison, strain accounts for 30% of total variance. Strain longevity combined across the two diets varies from 307 ± 37 days in BXD13 (*n* = 21) to 852 ± 33 days in BXD168 (*n* = 23). Some strains are fully resistant to the negative effects of HFD on body weight and life span while others are strongly affected, indicating substantial gene-by-diet interaction effects. While mean longevity on high fat is shortened by an average of 10%, genetic factors account for roughly two-fold range in life span. At least one strain, BXD8, actually has significantly improved longevity on the HFD (+37% on HF). Other strains such as BXD16 and BXD73, are immune to the high fat challenge with respect to change in life span. Of course, longevity of most strains is adversely affected (n = 67) and in the case of BXD65 life span is cut in half. Weight gain is characterized by a similar GXE effect—at least four family members are resistant to weight gain, including BXD16, BXD77, BXD87, and BXD91, gaining at most 5% over 100 days.

Our findings can be compared to strain variation and GXE effects in response to dietary restriction. Dietary restriction without malnutrition is regarded as having an almost universal benefit on longevity (Mair and Dillin, 2008; Masoro, 2009; Weindruch et al., 1986). One exception is a pair of studies on the impact of moderately intense restriction—a 40% reduction in caloric intake—across a large family of LXS strains of mice ((Liao et al., 2010; Rikke et al., 2010); *n* of up 44 strains with 10–20 replicates per strain). The most notable finding is the remarkably high variation in strain-specific change in longevity in response to dietary restriction. Life span is shortened in some LXS strains (maximum of 671-day loss), but lengthened in others (minimum of 300-day gain; GeneNetwork LXS phenotype 10164). Both the Liao and Rikke papers generated substantial controversy (Mattson, 2010) which was mirrored to some extent in matched studies of non-human primates (Mattison et al., 2017). Given such contrast in outcomes, it would be worthwhile extending the analysis of HFD to other dietary interventions in the expanded BXD family (Ashbrook et al., 2019).

### Diet and Morbidity

Major morbidities and likely causes of death among different members of the BXD family appear to be influenced by diet. Those on HFD have an increased prevalence and severity of cardiovascular disease and lesions. However, the effects of a very high fat diet on cardiovascular disease incidence is quite modest in mice, unlike that observed in human cohorts (Menotti and Puddu, 2015; Sacks Frank M. et al., 2017). Interestingly, incidence of sarcomas and carcinomas are higher on the CD than HFD. Large prospective studies have failed to detect strong associations between dietary fat and cancer risk (Willett, 2000). Evaluating a causal role of diet-induced obesity in the etiology of several chronic diseases and cancers has been difficult due to correlations with numerous lifestyle factors and resulting confounding biases. A growing body of literature indicates that interactions between select adipose tissue components and adjacent developing cancers are impacted during progression to obesity (Cozzo et al., 2017). Human epidemiology is increasingly turning to Mendelian randomization to assess the possible causal associations between risk factors and diseases (Gao et al., 2016). Well-controlled animal experiments could similarly provide additional understanding of causal associations and mechanisms underlying such complex relations.

### GXE, Weight Gain, and Associations with Longevity

Chronically high levels of fat consumption lead to substantial weight gain, associated metabolic disorders, and shortened lifespan, but causal interactions among these critical traits remains controversial. For example, studies exploiting Mendelian randomization have not shown a compelling link of triglyceride levels on longevity in humans (Liu et al., 2017). In general, a diet high in fat leads to obesity and reduced lifespan in diverse species including *Drosophila, C. elegans*, and mice (Otabe et al., 2007, Yen and Curran, 2016, Gáliková and Klepsatel, 2018). In our study, higher youthful body weight—between 100 to 200 days of age—is associated with a reduction in longevity of 4 to 5 days per gram. This corroborates much previous work that also demonstrates that larger body size within species is associated with shorter life span. For example, in outcrossed mice, for each gram increase in weight at 180 days, longevity is reduced by 10 to 15 days (Miller et al., 2000, their Figure 1). Among breeds of dog, for each kilogram increase in weight, longevity is reduced by ∼15 days (Kraus et al., 2013). In young adult and middle-aged humans, for each kilogram increase in body weight, longevity is reduced by 80 to 146 days (Samaras et al., 2002, their figure 6). In these three species, a 5% gain in young adult body weight is associated with a 1–3% loss in longevity. In both mice and dogs the relation between early weight gain and longevity is in part a well-known function of growth hormone (GH)–insulin-like growth factor 1 (IGF1) activity, although other mechanisms and gene variants are also likely to contribute. In humans, causes of this relation are probably a result of an interplay of many factors, especially nutritional composition, smoking, mean activity levels, and health care systems. For example, tall stature protects against cardiovascular disease due to lower levels of adiposity, lipid fractions, blood pressure, and better lung function (Nüesch et al., 2016). The linkage of longevity to IGF1 levels is also more tenuous in humans than in mouse and dog (Sanders et al., 2018), although short stature and lower GH/IGF1 signaling is associated with resistance to most types of cancers (Choi et al., 2019; Nunney, 2018).

In humans an increase in BMI from 27 to 42 kg/m^2^ increases all-cause mortality hazard ratio 1.6-fold (Wade et al., 2018). We see an even greater effect in the BXD family—the HFD diet increases weight 1.8-fold and the mortality hazard ratio 2-fold. But these average hazard ratios mask impressive modulation by genetic differences. While 45 of 57 family members gain significant weight, the other 12 gain only modest and statistically insignificant weight. For example, even after 400 days on the HFD, BXD16 is only 1.05-fold heavier than control, whereas BXD24 is 2.1-fold heavier. Remarkably, only 10% of the effect of diet on longevity is mediated through weight gain. HFD itself exerts a stronger direct effect on longevity in female BXDs than weight gain *per se*.

### Future Directions

Impressive GXE differences among the BXD family emphasize the complexity of interactions among diet, weight gain, and longevity. These effects need to be teased apart and explained at genetic, epigenetic, and mechanistic levels—work that is now in progress using the same strains and cohorts (Andreux et al., 2012; Houtkooper et al., 2013; Jha et al., 2018a, 2018b; Sandoval-Sierra et al., 2019; Williams et al., 2016; Wu et al., 2014). Substantial diversity in outcomes among close-knit members the BXD family highlight the fact that population averages obscure major GXE effects. Not much can be said with certainty from a single genometype or even a small number of diverse strains. In our case, detecting strong GXE interactions required data from 10 isogenic cases each of more than 67 strains under both conditions. Data of this type can be a foundation for specific predictions and to design a second wave of interventional studies to extend longevity and vigor (Miller et al., 2007; Strong et al., 2013). When extrapolated to humans, our results indicate that conventional dietary recommendations will be far too generalized to provide effective individual guidance. We emphasize that genotype and GXE effects are far stronger modulators of longevity and weight gain than average population effects.

## Acknowledgements

We thank Dr. James F. Nelson for helpful discussion on the LXS dietary restriction datasets. Thanks to David Ashbrook for proof-reading our final version of the manuscript.

## Author contributions

Conceptualization: EGW, JI, JA, LL, RWW

Aging Colony Management: SR, JI, CJ, MM

Investigation: SR, JI, CJ, MM, AC, KM, MM, WZ, JH, SN, LW, TS, CK, LL, RWW

Formal Analysis and Data Curation: SR, MBS, PJ, EGW, AS, MH, RWR, SS, RWW

Writing-Original Draft: SR, RWW

Writing-Review and Editing: SR, MBS, EGW, LM, RAM, JA, RWW

Companion Web Resources: AC, SR, RWW

## References

Andreux, P.A., Williams, E.G., Koutnikova, H., Houtkooper, R.H., Champy, M.-F., Henry, H., Schoonjans, K., Williams, R.W., and Auwerx, J. (2012). Systems Genetics of Metabolism: The Use of the BXD Murine Reference Panel for Multiscalar Integration of Traits. Cell 150, 1287–1299.

Ashbrook, D.G., Arends, D., Prins, P., Mulligan, M.K., Roy, S., Williams, E.G., Lutz, C.M., Valenzuela, A., Bohl, C.J., Ingels, J.F., et al. (2019). The expanded BXD family of mice: A cohort for experimental systems genetics and precision medicine. BioRxiv 672097.

Austad, S.N., and Fischer, K.E. (2016). Sex Differences in Lifespan. Cell Metab. 23, 1022–1033.

Barrington, W.T., Wulfridge, P., Wells, A.E., Rojas, C.M., Howe, S.Y.F., Perry, A., Hua, K., Pellizzon, M.A., Hansen, K.D., Voy, B.H., et al. (2018). Improving Metabolic Health Through Precision Dietetics in Mice. Genetics 208, 399–417.

Belknap, J.K. (1998). Effect of within-strain sample size on QTL detection and mapping using recombinant inbred mouse strains. Behav. Genet. 28, 29–38.

Cheng, C.J., Gelfond, J.A.L., Strong, R., and Nelson, J.F. (2019). Genetically heterogeneous mice exhibit a female survival advantage that is age- and site-specific: Results from a large multi-site study. Aging Cell 18, e12905.

Choi, Y.J., Lee, D.H., Han, K.-D., Yoon, H., Shin, C.M., Park, Y.S., and Kim, N. (2019). Adult height in relation to risk of cancer in a cohort of 22,809,722 Korean adults. Br. J. Cancer 120, 668–674.

Cozzo, A.J., Fuller, A.M., and Makowski, L. (2017). Contribution of Adipose Tissue to Development of Cancer. Compr. Physiol. 8, 237–282.

De Haan, G., and Van Zant, G. (1999). Genetic analysis of hemopoietic cell cycling in mice suggests its involvement in organismal life span. FASEB J. Off. Publ. Fed. Am. Soc. Exp. Biol. 13, 707–713.

Finkel, T. (2015). The metabolic regulation of aging. Nat. Med. 21, 1416–1423.

Fontana, L., and Partridge, L. (2015). Promoting Health and Longevity through Diet: From Model Organisms to Humans. Cell 161, 106–118.

Flurkey K, Currer JM, Harrison DE (2007). The mouse in biomedical research (Amsterdam; Boston: Elsevier, AP).

Gáliková, M., and Klepsatel, P. (2018). Obesity and Aging in the Drosophila Model. Int. J. Mol. Sci. 19, 1896.

Gao, C., Patel, C.J., Michailidou, K., Peters, U., Gong, J., Schildkraut, J., Schumacher, F.R., Zheng, W., Boffetta, P., Stucker, I., et al. (2016). Mendelian randomization study of adiposity-related traits and risk of breast, ovarian, prostate, lung and colorectal cancer. Int. J. Epidemiol. 45, 896–908.

Gelman, R., Watson, A., Bronson, R., and Yunis, E. (1988). Murine chromosomal regions correlated with longevity. Genetics 118, 693–704.

Griffiths, A.J., Miller, J.H., Suzuki, D.T., Lewontin, R.C., and Gelbart, W.M. (2000). Norm of reaction and phenotypic distribution. Introd. Genet. Anal. 7th Ed.

Hall, R.A., Liebe, R., Hochrath, K., Kazakov, A., Alberts, R., Laufs, U., Böhm, M., Fischer, H.-P., Williams, R.W., Schughart, K., et al. (2014). Systems Genetics of Liver Fibrosis: Identification of Fibrogenic and Expression Quantitative Trait Loci in the BXD Murine Reference Population. PLoS ONE 9.

Heilbronn, L.K., and Ravussin, E. (2003). Calorie restriction and aging: review of the literature and implications for studies in humans. Am. J. Clin. Nutr. 78, 361–369.

Hook, M., Roy, S., Williams, E.G., Bou Sleiman, M., Mozhui, K., Nelson, J.F., Lu, L., Auwerx, J., and Williams, R.W. (2018). Genetic cartography of longevity in humans and mice: Current landscape and horizons. Biochim. Biophys. Acta BBA - Mol. Basis Dis. 1864, 2718–2732.

Houtkooper, R.H., Argmann, C., Houten, S.M., Cantó, C., Jeninga, E.H., Andreux, P.A., Thomas, C., Doenlen, R., Schoonjans, K., and Auwerx, J. (2011). The metabolic footprint of aging in mice. Sci. Rep. 1.

Houtkooper, R.H., Mouchiroud, L., Ryu, D., Moullan, N., Katsyuba, E., Knott, G., Williams, R.W., and Auwerx, J. (2013). Mitonuclear protein imbalance as a conserved longevity mechanism. Nature 497, 451–457.

Jha, P., McDevitt, M.T., Halilbasic, E., Williams, E.G., Quiros, P.M., Gariani, K., Sleiman, M.B., Gupta, R., Ulbrich, A., Jochem, A., et al. (2018a). Genetic Regulation of Plasma Lipid Species and Their Association with Metabolic Phenotypes. Cell Syst. 6, 709-721.e6.

Jha, P., McDevitt, M.T., Gupta, R., Quiros, P.M., Williams, E.G., Gariani, K., Sleiman, M.B., Diserens, L., Jochem, A., Ulbrich, A., et al. (2018b). Systems Analyses Reveal Physiological Roles and Genetic Regulators of Liver Lipid Species. Cell Syst. 6, 722-733.e6.

Keipert, S., Voigt, A., and Klaus, S. (2011). Dietary effects on body composition, glucose metabolism, and longevity are modulated by skeletal muscle mitochondrial uncoupling in mice. Aging Cell 10, 122–136.

Kuningas, M., Mooijaart, S.P., Van Heemst, D., Zwaan, B.J., Slagboom, P.E., and Westendorp, R.G.J. (2008). Genes encoding longevity: from model organisms to humans. Aging Cell 7, 270–280.

Lang, D.H., Gerhard, G.S., Griffith, J.W., Vogler, G.P., Vandenbergh, D.J., Blizard, D.A., Stout, J.T., Lakoski, J.M., and McClearn, G.E. (2010). Quantitative trait loci (QTL) analysis of longevity in C57BL/6J by DBA/2J (BXD) recombinant inbred mice. Aging Clin. Exp. Res. 22, 8–19.

Liang, Y., Liu, C., Lu, M., Dong, Q., Wang, Z., Wang, Z., Xiong, W., Zhang, N., Zhou, J., Liu, Q., et al. (2018). Calorie restriction is the most reasonable anti-ageing intervention: a meta-analysis of survival curves. Sci. Rep. 8, 5779.

Liao, C.-Y., Rikke, B.A., Johnson, T.E., Diaz, V., and Nelson, J.F. (2010). Genetic Variation in the Murine Lifespan Response to Dietary Restriction: from Life Extension to Life Shortening. Aging Cell 9, 92–95.

Liu, Z., Burgess, S., Wang, Z., Deng, W., Chu, X., Cai, J., Zhu, Y., Shi, J., Xie, X., Wang, Y., et al. (2017). Associations of triglyceride levels with longevity and frailty: A Mendelian randomization analysis. Sci. Rep. 7, 41579.

de Magalhães, J.P., Wuttke, D., Wood, S.H., Plank, M., and Vora, C. (2012). Genome-Environment Interactions That Modulate Aging: Powerful Targets for Drug Discovery. Pharmacol. Rev. 64, 88–101.

Mattison, J.A., Colman, R.J., Beasley, T.M., Allison, D.B., Kemnitz, J.W., Roth, G.S., Ingram, D.K., Weindruch, R., de Cabo, R., and Anderson, R.M. (2017). Caloric restriction improves health and survival of rhesus monkeys. Nat. Commun. 8, 14063.

Mattson, M.P. (2010). Genes and behavior interact to determine mortality in mice when food is scarce and competition fierce. Aging Cell 9, 448–449.

McDaid, A.F., Joshi, P.K., Porcu, E., Komljenovic, A., Li, H., Sorrentino, V., Litovchenko, M., Bevers, R.P.J., Rüeger, S., Reymond, A., et al. (2017). Bayesian association scan reveals loci associated with human lifespan and linked biomarkers. Nat. Commun. 8.

Menotti, A., and Puddu, P.E. (2015). How the Seven Countries Study contributed to the definition and development of the Mediterranean diet concept: A 50-year journey. Nutr. Metab. Cardiovasc. Dis. 25, 245–252.

Miller, R.A., Chrisp, C., and Atchley, W. (2000). Differential longevity in mouse stocks selected for early life growth trajectory. J. Gerontol. A. Biol. Sci. Med. Sci. 55, B455–461.

Miller, R.A., Harper, J.M., Galecki, A., and Burke, D.T. (2002). Big mice die young: early life body weight predicts longevity in genetically heterogeneous mice. Aging Cell 1, 22–29.

Miller, R.A., Harrison, D.E., Astle, C.M., Floyd, R.A., Flurkey, K., Hensley, K.L., Javors, M.A., Leeuwenburgh, C., Nelson, J.F., Ongini, E., et al. (2007). An Aging Interventions Testing Program: study design and interim report. Aging Cell 6, 565–575.

Mitchell, S.J., Matute, J.M., Scheibye-Knudsen, M., Fang, E., Aon, M., González-Reyes, J.A., Cortassa, S., Kaushik, S., Gonzalez-Freire, M., Patel, B., et al. (2016). Effects of Sex, Strain, and Energy Intake on Hallmarks of Aging in Mice. Cell Metab. 23, 1093–1112.

Nüesch, E., Dale, C., Palmer, T.M., White, J., Keating, B.J., van Iperen, E.P., Goel, A., Padmanabhan, S., Asselbergs, F.W., Verschuren, W., et al. (2016). Adult height, coronary heart disease and stroke: a multilocus Mendelian randomization meta-analysis. Int. J. Epidemiol. 45, 1927–1937.

Nunney, L. (2018). Size matters: height, cell number and a person’s risk of cancer. Proc. R. Soc. B Biol. Sci. 285, 20181743.

Otabe, S., Yuan, X., Fukutani, T., Wada, N., Hashinaga, T., Nakayama, H., Hirota, N., Kojima, M., and Yamada, K. (2007). Overexpression of human adiponectin in transgenic mice results in suppression of fat accumulation and prevention of premature death by high-calorie diet. Am. J. Physiol. - Endocrinol. Metab. 293, E210–E218.

Peirce, J.L., Lu, L., Gu, J., Silver, L.M., and Williams, R.W. (2004). A new set of BXD recombinant inbred lines from advanced intercross populations in mice. BMC Genet. 5, 7.

Pennacchio, L.A., and Rubin, E.M. (2003). Comparative genomic tools and databases: providing insights into the human genome. J. Clin. Invest. 111, 1099–1106.

Rikke, B.A., Liao, C.-Y., McQueen, M.B., Nelson, J.F., and Johnson, T.E. (2010). Genetic dissection of dietary restriction in mice supports the metabolic efficiency model of life extension. Exp. Gerontol. 45, 691–701.

Sacks Frank M., Lichtenstein Alice H., Wu Jason H.Y., Appel Lawrence J., Creager Mark A., Kris-Etherton Penny M., Miller Michael, Rimm Eric B., Rudel Lawrence L., Robinson Jennifer G., et al. (2017). Dietary Fats and Cardiovascular Disease: A Presidential Advisory From the American Heart Association. Circulation 136, e1–e23.

Samaras, T.T., Storms, L.H., and Elrick, H. (2002). Longevity, mortality and body weight. Ageing Res. Rev. 1, 673–691.

Sandoval-Sierra, J.V., Helbing, A.H.B., Williams, E.G., Ashbrook, D.G., Roy, S., Williams, R.W., and Mozhui, K. (2019). Influence of body weight at young adulthood on the epigenetic clock and lifespan in the BXD murine family. BioRxiv 791582.

Saul, M.C., Philip, V.M., Reinholdt, L.G., and Chesler, E.J. (2019). High-Diversity Mouse Populations for Complex Traits. Trends Genet. 35, 501–514.

Skorupa, D.A., Dervisefendic, A., Zwiener, J., and Pletcher, S.D. (2008). Dietary composition specifies consumption, obesity and lifespan in Drosophila melanogaster. Aging Cell 7, 478–490.

Speakman, J.R., Mitchell, S.E., and Mazidi, M. (2016). Calories or protein? The effect of dietary restriction on lifespan in rodents is explained by calories alone. Exp. Gerontol. 86, 28–38.

Strong, R., Miller, R.A., Astle, C.M., Baur, J.A., de Cabo, R., Fernandez, E., Guo, W., Javors, M., Kirkland, J.L., Nelson, J.F., et al. (2013). Evaluation of Resveratrol, Green Tea Extract, Curcumin, Oxaloacetic Acid, and Medium-Chain Triglyceride Oil on Life Span of Genetically Heterogeneous Mice. J. Gerontol. A. Biol. Sci. Med. Sci. 68, 6–16.

Therneau, T.M., and Grambsch, P.M. (2000). Modeling Survival Data: Extending the Cox Model (New York: Springer-Verlag).

Vaughan, K.L., Kaiser, T., Peaden, R., Anson, R.M., de Cabo, R., and Mattison, J.A. (2018). Caloric Restriction Study Design Limitations in Rodent and Nonhuman Primate Studies. J. Gerontol. Ser. A 73, 48–53.

Viechtbauer, W. (2010). Conducting Meta-Analyses in R with the metafor Package. J. Stat. Softw. 36, 1–48.

Wade, K.H., Carslake, D., Sattar, N., Davey Smith, G., and Timpson, N.J. (2018). BMI and Mortality in UK Biobank: Revised Estimates Using Mendelian Randomization. Obes. Silver Spring Md 26, 1796–1806.

Willett, W.C. (2000). Diet and Cancer. The Oncologist 5, 393–404.

Williams, R.W. (2006). Animal models in biomedical research: ethics, challenges and opportunities. Princ. Mol. Med. 2nd edition, 53–60.

Williams, R.W., and Williams, E.G. (2017). Resources for Systems Genetics. In Systems Genetics, K. Schughart, and R.W. Williams, eds. (New York, NY: Springer New York), pp. 3–29.

Williams, E.G., Mouchiroud, L., Frochaux, M., Pandey, A., Andreux, P.A., Deplancke, B., and Auwerx, J. (2014). An Evolutionarily Conserved Role for the Aryl Hydrocarbon Receptor in the Regulation of Movement. PLOS Genet. 10, e1004673.

Williams, E.G., Wu, Y., Jha, P., Dubuis, S., Blattmann, P., Argmann, C.A., Houten, S.M., Amariuta, T., Wolski, W., Zamboni, N., et al. (2016). Systems proteomics of liver mitochondria function. Science 352, aad0189.

Wu, Y., Williams, E.G., Dubuis, S., Mottis, A., Jovaisaite, V., Houten, S.M., Argmann, C.A., Faridi, P., Wolski, W., Kutalik, Z., et al. (2014). Multilayered Genetic and Omics Dissection of Mitochondrial Activity in a Mouse Reference Population. Cell 158, 1415–1430.

Yen, C.A., and Curran, S.P. (2016). Gene-diet interactions and aging in C. elegans. Exp. Gerontol. 86, 106–112.

Yuan, R., Peters, L.L., and Paigen, B. (2011). Mice as a Mammalian Model for Research on the Genetics of Aging. ILAR J. Natl. Res. Counc. Inst. Lab. Anim. Resour. 52, 4–15.

