## Supplemental Figures S1-S3 for "Gene-by-environmental modulation of longevity and weight gain in the murine BXD family"

Supplemental Information

Figure S1. Related to Figure 1 and Figure 3. (A) Mean longevity on the low fat chow diet (CD) in 73 BXD strains. (B) Mean longevity on the high fat diet (HFD) in 74 BXD strains. (C) Mean body weight on CD at 500 days of age in 59 BXD strains. (D) Mean body weight on HFD at 500 days of age in 59 BXD strains.

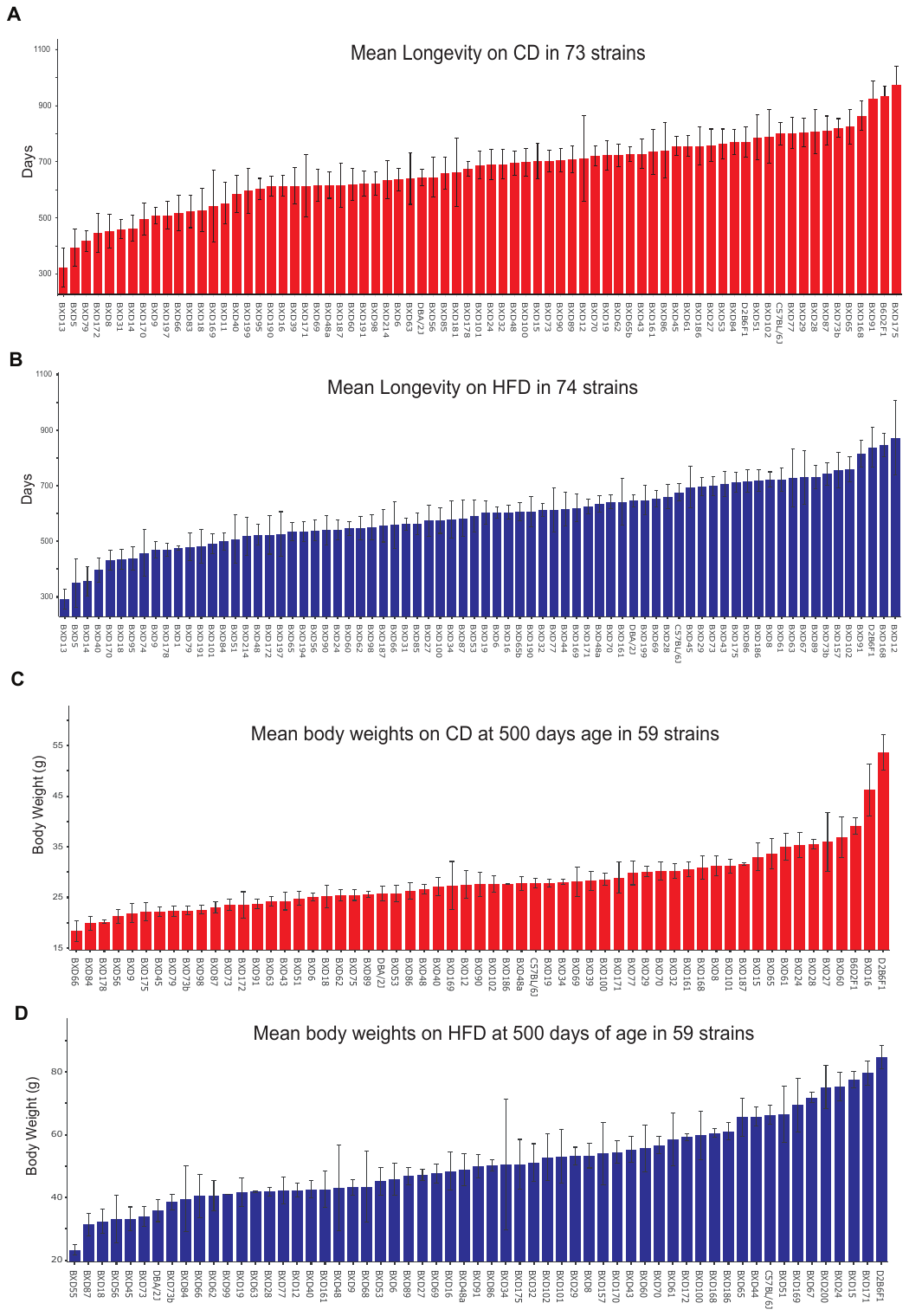

Figure S2. High fat diet does not modulate variability in longevity. Mean coefficients of variation (CV) for all strains (A) on CD with sample sizes of greater than 6 cases (*n* = 52, GN2 trait BXD_21533). Asterisk denotes the strain for which the CV value was winsorized. (B) on HFD with sample sizes of greater than 6 cases (*n* = 50, GN2 trait BXD_21534). (C) Correlation between CV on CD and HFD is not significant.

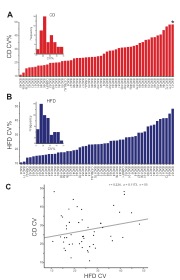

Figure S3. Association between longevity and major metabolic organ weights at ~500 days of age in a separate subsample of animals on the two diets. Red points represent individuals on CD and blue points those on HFD. The correlations between longevity and body and organ weights at 18 months in this subset of animals is not significant (A-F).

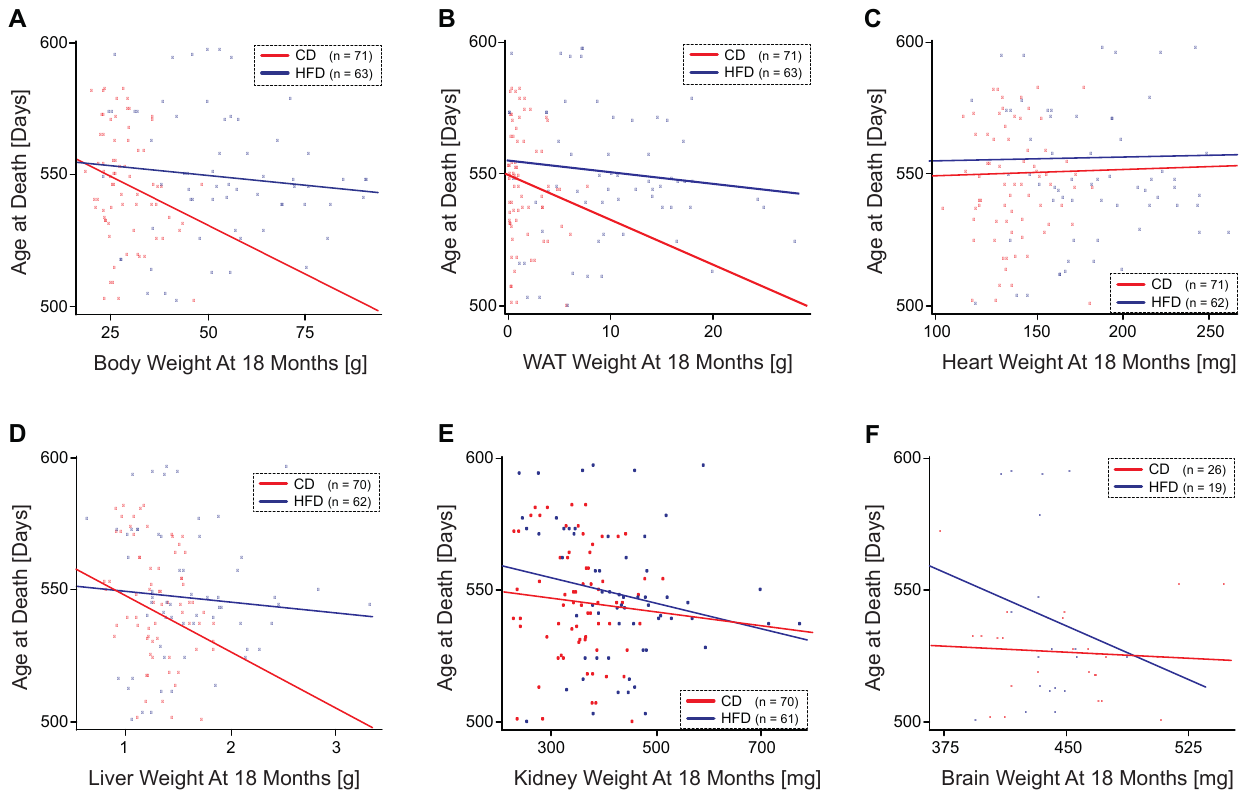
